# The secreted neuronal signal Spock1 regulates the blood-brain barrier

**DOI:** 10.1101/2021.10.13.464312

**Authors:** Natasha M. O’Brown, Nikit B. Patel, Ursula Hartmann, Allon M. Klein, Chenghua Gu, Sean G. Megason

**Author notes:** denotes corresponding author Correspondence: Natasha M. O’Brown Department of Systems Biology, Harvard Medical School Boston, MA 02115, U.S.A. Sean G. Megason Department of Systems Biology, Harvard Medical School Boston, MA 02115, U.S.A.

## Abstract

The blood-brain barrier (BBB) is a unique set of properties of the brain vasculature which severely restricts its permeability to proteins and small molecules. Classic chick-quail chimera studies showed that these properties are not intrinsic to the brain vasculature but rather are induced by surrounding neural tissue. Here we identify Spock1 as a candidate neuronal signal for regulating BBB permeability in zebrafish and mice. Mosaic genetic analysis shows that neuronally-expressed Spock1 is cell non-autonomously required for a functional BBB. Leakage in *spock1* mutants is associated with altered extracellular matrix (ECM), increased endothelial transcytosis, and altered pericyte-endothelial interactions. Furthermore, a single dose of recombinant SPOCK1 partially restores BBB function in *spock1* mutants by quenching gelatinase activity and restoring vascular expression of BBB genes including *mcamb*. These analyses support a model in which neuronally secreted Spock1 induces BBB properties by altering the ECM, thereby regulating pericyte-endothelial interactions and downstream vascular gene expression.

**One-Sentence Summary:** Spock1 is a signal secreted by neurons that induces barrier properties in the brain vasculature

## INTRODUCTION

The blood-brain barrier (BBB) maintains a tightly controlled homeostatic environment in the brain that is required for proper neural function. BBB breakdown has been implicated in multiple neurodegenerative diseases including Alzheimer’s, Parkinson’s, and Huntington’s Diseases (Sweeney et al., 2018). Conversely, the BBB also serves as an obstacle for effective drug delivery to the brain. Therefore there is great interest in generating a better understanding of how to therapeutically regulate its permeability, both to restore BBB function in neurodegeneration and to transiently open it for improved chemotherapeutic access for brain tumors. The BBB is a specialized property of the brain vasculature, which is composed of a thin, continuous layer of non-fenestrated endothelial cells with uniquely restrictive properties. Brain endothelial cells create the barrier via two primary cellular mechanisms: 1) specialized tight junction complexes that block the transit of small water-soluble molecules between cells and 2) reduced levels of vesicular trafficking or transcytosis to restrict transit through endothelial cells (Reese and Karnovsky, 1967). At a molecular level, functional barrier endothelial cells are differentiated from peripheral endothelial cells by high expression of the tight junction molecule Claudin-5 (Campbell et al., 2008) and the fatty acid transporter Mfsd2a (Andreone et al., 2017; Ben-Zvi et al., 2014), and an absence of the fenestra and vesicle associated protein Plvap (Hallmann et al., 1995). Expression of substrate-specific influx and efflux transporters, which dynamically regulate the intake of necessary nutrients and the removal of metabolic waste products (Sanchez-Covarrubias et al., 2014; Umans et al., 2017), further enhances the selectivity of the BBB.

The restrictive properties of the BBB are not intrinsic to brain endothelial cells, but are induced and maintained by signals in the brain microenvironment during embryonic development (Stewart and Wiley, 1981). When avascularized quail neural tissue was transplanted into embryonic chicken gut cavities, the blood vessels ingressing from the non-barrier gut tissue obtained functional BBB properties, specifically both tight junctions and reduced levels of vesicles, resulting in Trypan blue tracer confinement within the blood vessels (Stewart and Wiley, 1981). Conversely, when avascularized quail somite tissue was transplanted into embryonic chick brain, the blood vessels ingressing from the neural tissue lost functional BBB properties, leaking tracer into the quail graft due to increased endothelial transcytosis and loss of tight junction function. These data show that signals from the neural microenvironment to the vasculature control barrier function. Furthermore, these microenvironmental signals are also required to actively maintain barrier properties throughout life (Guérit et al., 2021; Heithoff et al., 2021; Lyck et al., 2009; Urich et al., 2012).

Mural cells called pericytes share the endothelial basement membrane and comigrate with endothelial cells during neovascularization of the developing brain. This close interaction between pericytes and endothelial cells is required for both the initiation and continued maintenance of barrier properties (Armulik et al., 2010; Bell et al., 2010; Daneman et al., 2010). Astrocytes, glial cells found exclusively in the central nervous system, completely ensheath the vasculature with their endfeet in a polarized fashion (Abbott et al., 2006). Astrocytes arise late in embryonic development, after neurogenesis is complete, with the vast majority of gliogenesis occurring postnatally in rodents. Mammalian astrocytes are potent inducers of barrier properties in vitro and in vivo, and are required for BBB maintenance (Guérit et al., 2021; Heithoff et al., 2021; Janzer and Raff, 1987). While the necessity of these two cell types has been known for decades, canonical Wnt signaling arising from both neuronal and astrocytic sources is the only microenvironmental signal known to induce and maintain BBB function (Benz et al., 2019; Daneman et al., 2009; Guérit et al., 2021; Liebner et al., 2008; Martin et al., 2022; Stenman et al., 2008; Vanhollebeke et al., 2015; Wang et al., 2012, 2018). However in addition to its role in regulating barrier properties, Wnt signaling also plays an important role in tip cell specification and angiogenesis in the developing brain (Stenman et al., 2008; Vanhollebeke et al., 2015; Wang et al., 2012). Furthermore, mouse endothelial cells are Wnt-responsive as early as E9.5 (Liebner et al., 2008), preceding the acquisition of functional barrier properties (Daneman et al., 2009), suggesting that other signaling may be acting in conjunction with or in addition to Wnt signaling for the induction of BBB properties.

To further examine the molecular determinants of BBB development, we turned to the optically transparent zebrafish system, which allows for imaging of the intact BBB in toto throughout zebrafish development. Zebrafish brain endothelial cells express many of the same molecular markers as mammalian brain endothelial cells, including Glut1, Cldn5, ZO-1, and Mfsd2a (Guemez-Gamboa et al., 2015; Jeong et al., 2008; O’Brown et al., 2019; Umans et al., 2017; Vanhollebeke et al., 2015; Xie et al., 2010). We previously characterized the molecular and subcellular mechanisms of functional BBB development in zebrafish, and determined that the zebrafish BBB becomes functionally mature by 5 days post fertilization (dpf) due to the suppression of transcytosis rather than through the acquisition of tight junction function, which was observed as early as 3 dpf (O’Brown et al., 2019). During these studies, we serendipitously discovered a recessive viable mutant with profound forebrain and midbrain barrier leakage which we identify here to be the neuronally produced and secreted proteoglycan Spock1. Using an array of perturbations and imaging techniques, we uncover the range of Spock1 signaling to the vasculature, and identify that mechanistically Spock1 signaling acts through control of the pericyte-endothelial extracellular matrix (ECM) to control endothelial transcytosis.

## RESULTS

### Identification and mapping of a spontaneous leaky mutant to *spock1*

We found a spontaneous recessive mutant that leaks both an injected 1 kDa Alexa Fluor (AF) 405 NHS Ester dye and a transgenic 80 kDa serum protein DBP-EGFP (Xie et al., 2010) into the forebrain and midbrain as early as 3 days post fertilization (dpf) with no improvement in BBB function throughout larval development (Figure 1A-E) or adulthood (Figure 1G). This increased BBB permeability in homozygous mutants occurs in the absence of hemorrhage or vascular patterning defects (Figure S1), or any reduction in viability or fertility. Time lapse imaging of injected 10 kDa Dextran revealed a steady accumulation of Dextran in the brain parenchyma of leaky mutants over the course of an hour (Figure S2), closely resembling the dynamics of Dextran leakage observed in *mfsd2aa* mutants with increased endothelial transcytosis (O’Brown et al., 2019).

**Figure 1.**
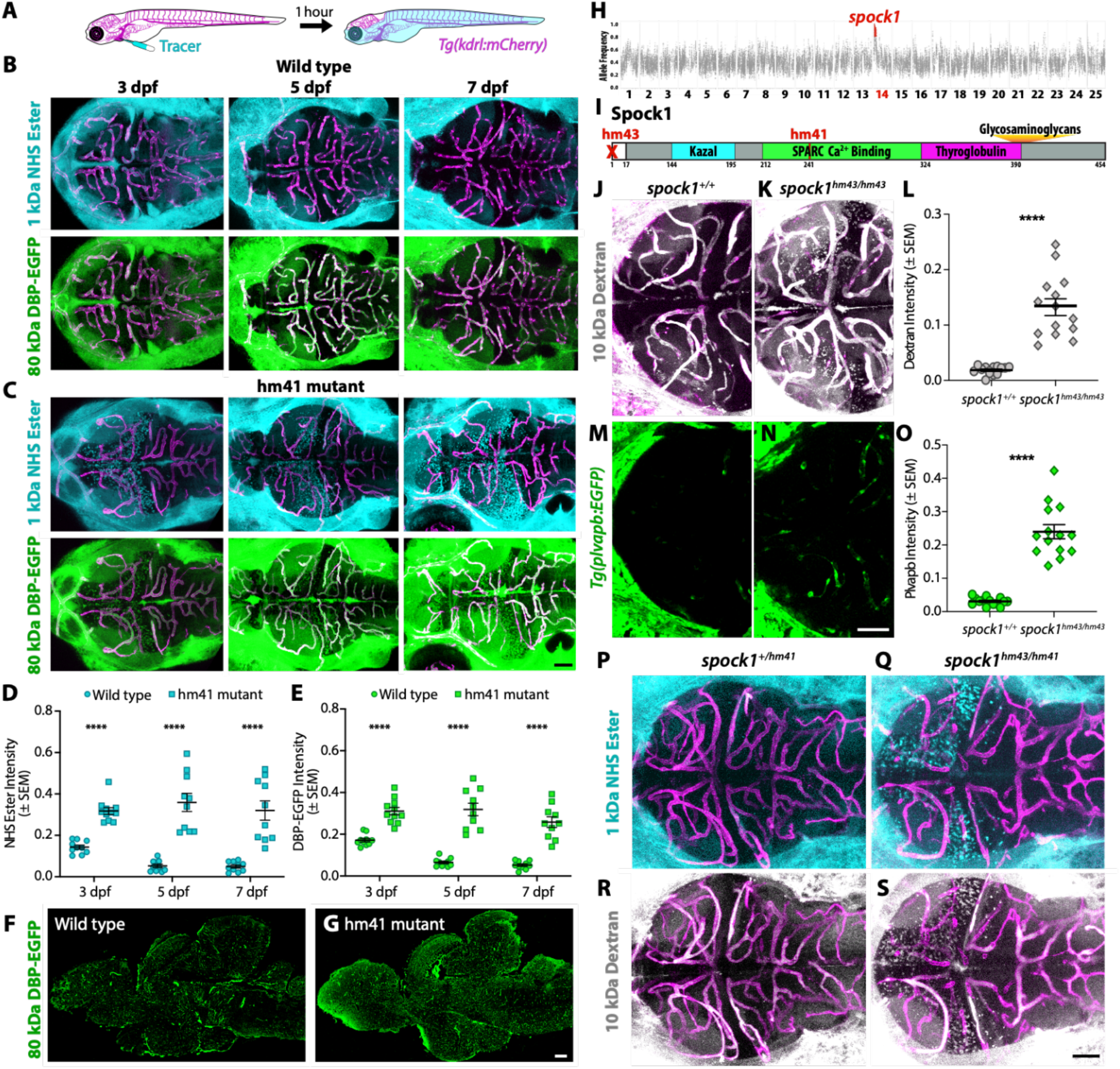
Spock1 induces BBB functional development. (**A-E**) Fluorescent tracer leakage assays in larval zebrafish reveal BBB leakage in the forebrain and midbrain of both injected 1 kDa Alexa Fluor 405 NHS Ester (turquoise) and 80 kDa DBP-EGFP (green) of *hm41* homozygous larvae (C) compared to wild type controls (B) throughout larval development, as quantified in D and E. (**F-G**) Sagittal sections of adult brains show that the mutant leakage persists into adulthood (G). (**H**) Linkage mapping of the *hm41* mutant phenotype reveals tight linkage to *spock1* on chromosome 14. (**I**) *Spock1^hm41^* has several point mutations (T241A and several silent mutations) in the SPARC domain. *Spock1^hm43^* has a deletion of the 5’ UTR and start codon. (**J-O**) Dextran leakage assays show *spock1^hm43/hm43^* mutants (K) have increased BBB leakage (L). *spock1^hm43/hm43^* mutants have increased expression of the leaky vessel marker plvapb (N and O). (**P-S**) Compound *spock1^hm43/hm41^* heterozygotes also display increased NHS Ester (Q) and Dextran (S) leakage compared to *spock1^hm41/+^* heterozygote siblings, which confine both injected tracers at 5 dpf (P and R). Scale bars represent 50 µm (C, N, S) and 200 µm (G). **** p<0.0001 by 2way ANOVA (D and E) and by t test (L and O).

To identify the mutation responsible for this leakage phenotype, we performed linkage mapping on 5 dpf mutant and wild type siblings using bulk segregant RNAseq (Miller et al., 2013) and identified a single peak at chr14:2205271-3513919 (GRCz11; Figures 1H and S3). Of the 14 genes within the linkage region (Figure S3), 8 were expressed at 5 dpf. Two of these genes were differentially expressed, *csf1ra* and *gstp2* (Table S1), and one had several mutations that segregated with the leakage phenotype, *spock1.* To test whether loss of any of these genes conferred the increased BBB permeability, we assessed tracer leakage in mosaic crispants (zebrafish larvae injected at the 1-cell stage with Cas9 protein and gene-specific sgRNAs) at 5 dpf and observed no BBB defects in 42 *gstp2* or 35 *csf1ra* crispants (data not shown). However, when we assessed BBB function in *spock1* crispants, we observed a strong leaky phenotype in 26% of larvae (26/100 injected fish; Figure S3) and moderate leakage in 53% (53/100 injected fish; Figure S3) as well as a few that displayed BBB leakage restricted to one midbrain hemisphere (4/100 injected fish; Figure S3). Leakage in *spock1* crispants corresponded with a loss of *spock1* expression (Figure S3). *Spock1* encodes a secreted protein of unclear function named for its conserved protein domains: <u>SP</u>ARC (<u>O</u>steonectin), <u>C</u>wcv And <u>K</u>azal Like Domains Proteoglycan <u>1</u> protein (also known as Testican-1). Spock1 has three predicted domains: 1) a Kazal-type serine protease inhibitor, 2) an extracellular SPARC calcium-binding region, and 3) a thyroglobulin type-1 repeat region, in addition to being decorated by both chondroitin sulfate (CS) and heparan sulfate (HS) glycosaminoglycan (GAG) chains (Figure 1I) (Bonnet et al., 1996; Edgell et al., 2004). Further sequencing of *spock1* confirmed a few point mutations in the SPARC calcium-binding domain in the leaky fish (Figure 1I), including a T241A missense mutation and two silent mutations at Q249 and G250, hereafter referred to as s*pock1^hm41/hm41^* mutants.

To validate that loss of Spock1 function causes the leakage phenotype in the spontaneous mutants, we generated a stable null allele designated *spock1^hm43^* using CRISPR mutagenesis with two sgRNAs to completely delete the 5’ UTR and start codon of *spock1* (Figure 1I). As expected based on the 550 bp genomic deletion, *spock1^hm43/hm43^*fish do not express *spock1* (Figure S4). Functional tracer leakage assays using 10 kDa Dextran revealed the same increased BBB permeability specifically in the forebrain and the midbrain observed in the spontaneous s*pock1^hm41/hm41^* mutants (Figure 1J-L). This increased BBB permeability was accompanied with increased expression of the pathological marker of leaky endothelial cells *plvapb:EGFP* (Umans et al., 2017) compared to the negligible levels observed in wild type siblings with a functional BBB (Figure 1M-O). Lastly, we crossed both lines to test for genetic complementation, and observed the leaky phenotype in *spock1^hm41/hm43^* compound heterozygotes (Figure 1P-S), further indicating that the initial spontaneous mutant acts via loss of Spock1 function.

### Neuronal Spock1 signals to the vasculature within a short range

Given the necessity of Spock1 in determining BBB function, we next wanted to assess where *spock1* was expressed during BBB development. Using HCR fluorescent *in situ* hybridization, we determined that *spock1* mRNA is expressed throughout the developing central nervous system (CNS), including the entire brain, retina and spinal cord (Figures 2A and S4), and that this expression is unaltered in the spontaneous *spock1^hm41/hm41^* mutants (Figure S4). Closer examination of the fluorescent signal revealed co-localization with the neuronal marker *elavl3* and not the vascular marker *kdrl* (Figure 2D-G), similar to its predominantly neural expression in the developing mouse CNS (Charbonnier et al., 2000). Prior single-cell RNA sequencing (scRNA-seq) data (Raj et al., 2020) and our own also showed expression of *spock1* primarily in neurons and never in vascular endothelial cells (Figure S6). Taken together, these data indicate that we have identified a neuronal signal, Spock1, that plays a role in establishing endothelial BBB properties during development without altering vascular patterning.

**Figure 2.**
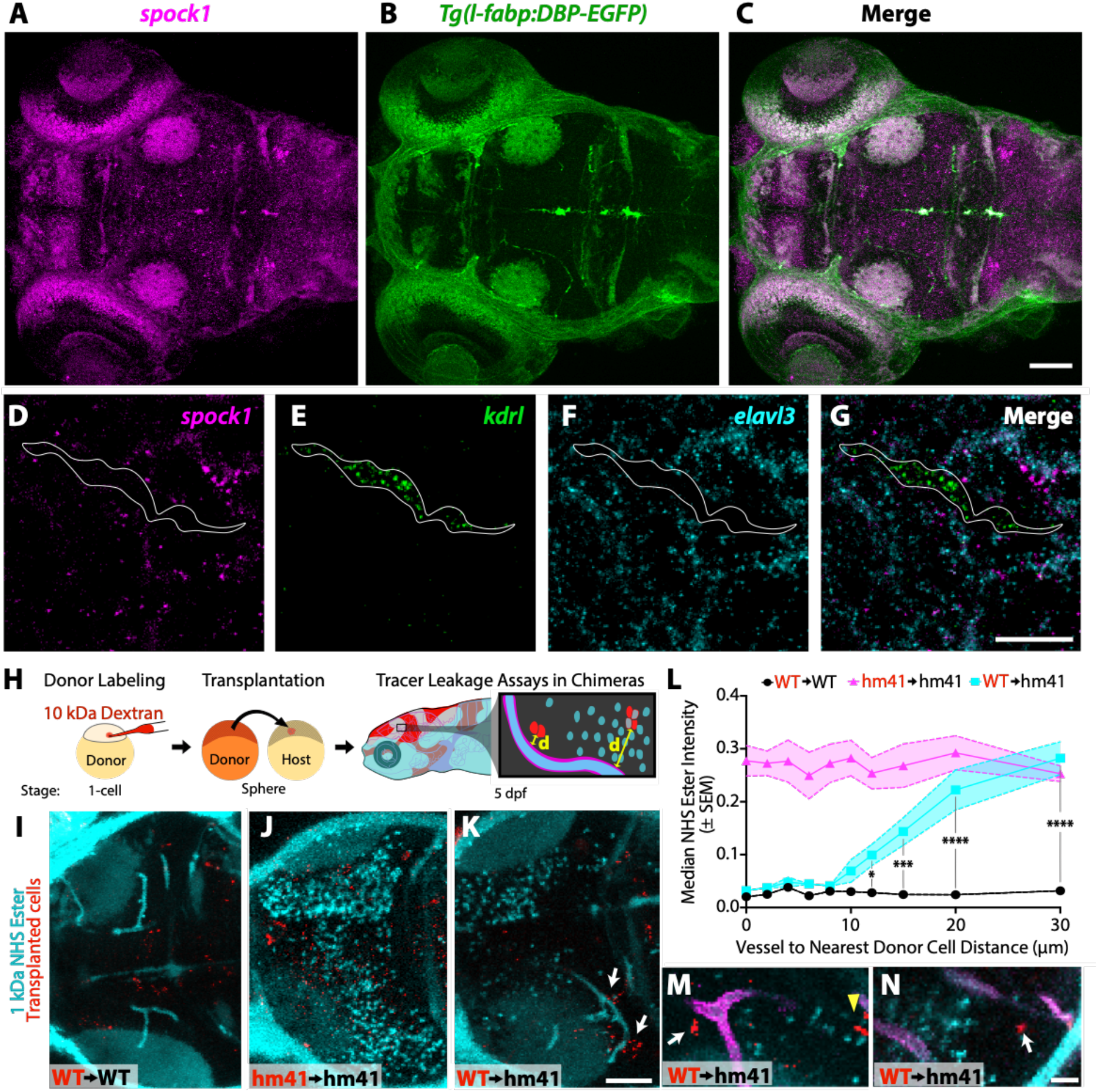
Neuronal Spock1 regulates endothelial cells within a 10-20 µm range. (**A-C**) Whole-mount HCR *in situ* hybridization for *spock1* (A) reveals signal throughout the entire 5 dpf brain, which is outlined by the fixed DBP-EGFP serum transgene (B). (**D-G**) Slide HCR *in situ* hybridization for *spock1* (D), the vascular transcript *kdrl* (E), and the neuronal transcript *elavl3* (F) reveal the lack of *spock1* signal in the vasculature (outlined in white). (**H**) Schematic of transplantation experiments. Cells from donor embryos labeled with 10 kDa Dextran (red) at the single cell stage are transplanted into unlabeled host embryos at sphere stage. Leakage of the injected 1 kDa AF 405 NHS Ester (turquoise) into the midbrain parenchyma is then measured in relationship to the distance (d) from the blood vessel to the nearest donor cell in the 5 dpf chimeric larvae. (**I-K**) Representative dorsal 20 µm thick maximum intensity projection confocal image of a chimeric larva with transplanted wild type donor cells (red) into a wild type host (I, WT→WT), *spock1^hm41/hm41^* donor cells into a *spock1^hm41/hm41^* mutant host (J, hm41→hm41), and wild type donor cells into a *spock1^hm41/hm41^* mutant host (K, WT→hm41). (**L**) Quantification of the NHS leakage in WT→WT (black line) and hm41→hm41 (magenta line) reveals no change in tracer leakage in relationship to the nearest donor cell, with wild type fish confining the tracer and mutant fish leaking the tracer. However WT→hm41 (turquoise line) transplants reveal a full rescue of the leakage in the mutant background when the transplanted cell is within 10 µm of a blood vessel and no effect if the donor cell is further than 20 µm from the vessel. (**M-N**) Zoomed in images of WT→hm41 transplants. The white arrows point to instances of local rescue of tracer (turquoise) leakage when the wild type donor cell is close but not directly contacting the mutant vasculature (magenta). The yellow arrowhead marks wild type donor cells that fall outside the range of Spock1 signaling. Scale bars represent 50 µm (D and K) and 10 µm (G and N). * p=0.0495, *** p=0.0001, **** p<0.0001 by 2way ANOVA compared to WT→WT transplants.

To determine the range of the Spock1 signal, we performed cell transplantation experiments to make genetically mosaic zebrafish embryos and assessed tracer leakage in relationship to the closest transplanted cells at 5 dpf (Figure 2H). When we transplanted labeled wild type cells into wild type host embryos, we observed a negligible level of tracer accumulation in the brain parenchyma regardless of the proximity of the nearest donor cell (Figure 2I and 2L). In contrast, when we transplanted *spock1^hm41/hm41^* mutant cells into mutant host embryos, we observed high levels of tracer accumulation in the parenchyma regardless of the proximity of the nearest donor cell (Figure 2J and 2L). Strikingly, when we transplanted wild type cells into *spock1* mutant hosts, we observed a complete rescue of the mutant leakage if the wild type donor cell was within 10 µm of a blood vessel and no rescue if the donor cell was more than 20 µm away (Figure 2K-N), indicating that Spock1 can functionally signal at a distance of 10 to 20 µm. Importantly, wild type cells could rescue leakage in mutant hosts when they differentiate as neurons but not endothelial cells (Figure 2M).

### Mutant leakage arises from increased transcytosis and altered pericyte-endothelial interactions

To assess the subcellular mechanism of increased BBB permeability in *spock1^hm41/hm41^* mutants, we injected electron-dense NHS-gold nanoparticles (5 nm) into circulation, followed by transmission electron microscopy (TEM) imaging in 7 dpf *spock1* mutant and wildtype siblings. These TEM leakage assays revealed no alterations in the cellular composition of the neurovascular unit (NVU) in *spock1* mutants, with endothelial cells and pericytes sharing a basement membrane that is surrounded by neurons and glia (Figure 3A and 3B). A modest impairment in tight junction function was observed in *spock1* mutants (52/59 functional tight junctions; Figure 3D) compared to wild type siblings (57/57; Figure 3C). However, *spock1* mutants had a significant increase in both small (<100 nm diameter) non-clathrin coated vesicles (Figure 3D and 3G) and large (>200 nm diameter) vesicles (Figure 3F and 3H) suggesting that the leakage results primarily from an increase in vesicular trafficking across endothelial cells. In addition to the increase in total large vesicular abundance in *spock1* mutants, we also observed several of these large vesicles fused to the abluminal membrane in multiple larvae with a few examples of the gold nanoparticles visibly spilling into the endothelial basement membrane (Figure 3F). Although pericyte coverage of the endothelium was unaltered (Figure S1), *spock1* mutants displayed overall thinner pericyte-endothelial basement membranes with several pericytes having long stretches of apparently direct contact on endothelial cells (Figure 3I-K), corresponding with observations by live confocal microscopy (Figure S1). These data suggest that loss of Spock1 function alters the critical pericyte extracellular interactions with endothelial cells, resulting in loss of BBB properties as observed in pericyte-deficient mice and zebrafish (Armulik et al., 2011; Bell et al., 2010; Daneman et al., 2010; Wang et al., 2013).

**Figure 3.**
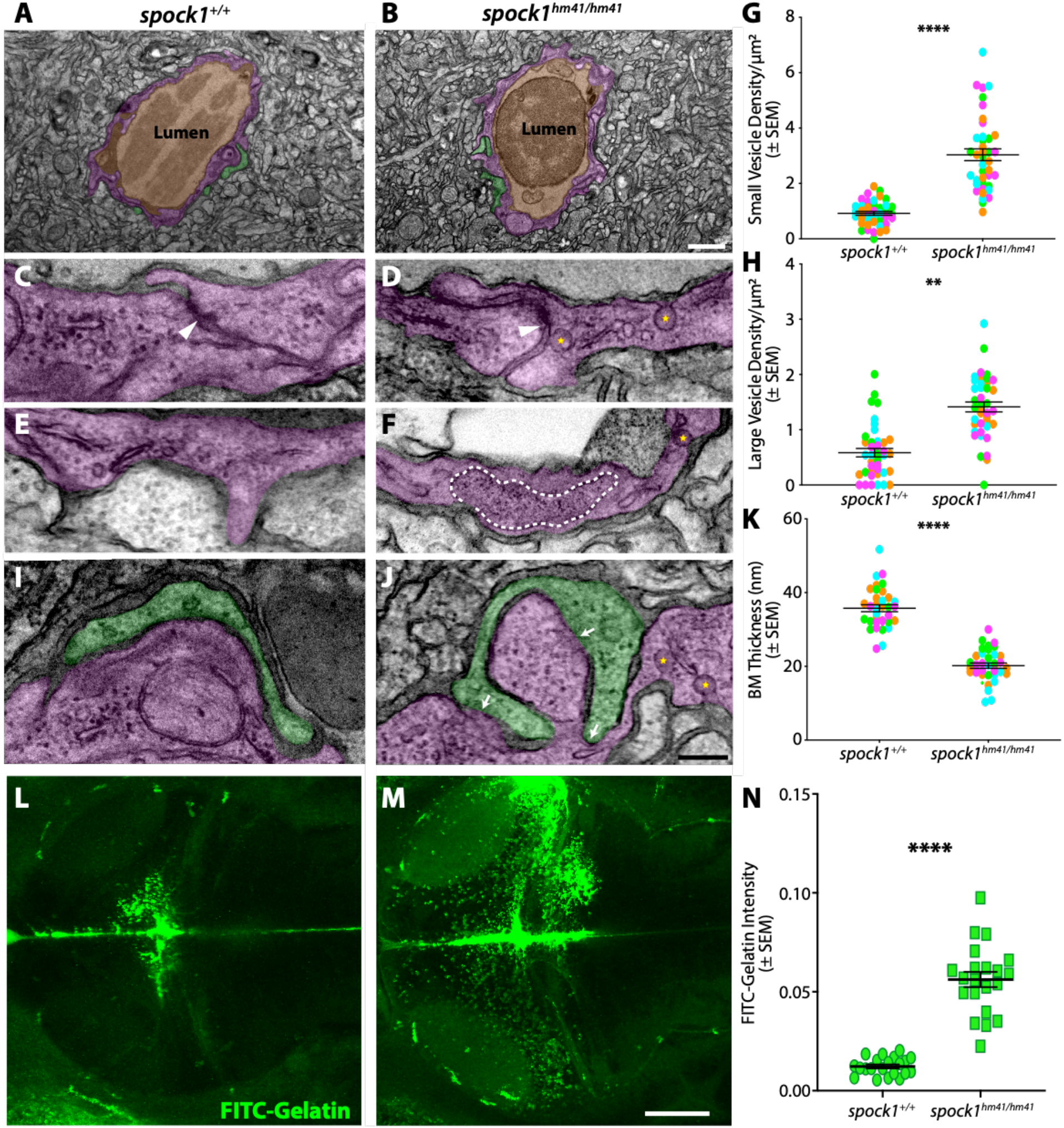
Spock1 mutants have increased endothelial vesicles. (**A-B**) The neurovascular unit remains intact in *spock1^hm41/hm41^* mutants (B) with a continuous single layer of endothelial cells (pseudocolored magenta) enclosing the lumen (pseudocolored orange) and in close contact with pericytes (pseudocolored green). (**C-H**) The majority of tight junctions (white arrowheads) are functionally restrictive in the *spock1^hm41^*mutant endothelial cells (88%). Mutant endothelial cells displayed a significant increase in vesicular density, including both small flask shaped vesicles (yellow stars, G) and larger vesicles greater than 200 nm in diameter (outlined by white dashed line in F, H). (**I-K**) While pericyte coverage is unaltered in *spock1^hm41^* mutants, the pericyte-endothelial cell interactions are altered in the mutants, with several instances of direct pericyte-endothelial cell contact (white arrows) and overall diminished average basement membrane thickness between the two vascular cells (K). (**L-N**) In vivo gelatin zymography in wild type (L) and *spock1^hm41/hm41^* mutants (M) reveals significantly increased gelatinase activity in the mutant midbrain compared to wild type siblings (N). Scale bars represent 1 µm (B), 200 nm (J), and 50 µm (M). ** p=0.0029 (H), **** p<0.0001 by nested t test (G and K) and by t test (N).

One of the proposed roles of Spock1 is to regulate matrix metalloproteinase (MMP) activation, specifically gelatinases MMP-2 and MMP-9 (Du et al., 2020; Edgell et al., 2004; Nakada et al., 2001; Váncza et al., 2022; Ye et al., 2020). To assess whether *spock1^hm41/hm41^* mutants had altered gelatinase activity that corresponded to observed changes in vascular cell interactions, we performed *in vivo* gelatin zymography by intracranial injections of highly quenched FITC-gelatin. Proteolytic cleavage of FITC-gelatin by either MMP-2 or 9 releases bright FITC peptides resulting in a non-reversible increase in fluorescence. This increase in fluorescence upon digestion is proportional to the proteolytic activity of the MMPs, allowing for the detection of gelatinase activity *in vivo* (Garcia-Alloza et al., 2009; Underly et al., 2017). Using this assay, we were able to see that in wild type 5 dpf fish, the majority of the signal appears strongly in the cerebral spinal fluid (CSF) and more subtly in many pericytes (Figure 3L). While *spock1^hm41/hm41^* mutants also displayed strong gelatinase activity in the CSF, this was accompanied by a regional increase in gelatinase activity specifically in the forebrain and midbrain (Figure 3M and 3N), the same regions that were also prone to tracer accumulation (Figure 1). These data suggest that Spock1 helps induce barrier function by keeping gelatinase activity in check.

### Spock1 plays a conserved role in inducing barrier function in mammals

To determine if Spock1 plays a conserved role in determining vertebrate BBB properties, we assessed BBB function in embryonic day 15.5 (E15.5) *Spock1^-/-^* mice (Röll et al., 2006) (Figure 4A), a stage when the cortex BBB is fully functional (Ben-Zvi et al., 2014). While wild type siblings confined both 550 Da Sulfo-NHS-Biotin (Figure 4B) and 10 kDa Dextran (Figure 4D) within the brain vasculature, *Spock1^-/-^* mice leaked both tracers into the cortex parenchyma (Figure 4C-G). This increased BBB permeability was also associated with increased expression of the PLVAP (Figure 4H-L), as seen in the zebrafish mutants (Figure 1N). These data indicate the Spock1 plays a conserved role in inducing barrier properties during embryonic development. However, when we repeated these leakage assays in adult mice, we observed full recovery in BBB function in *Spock1^-/-^* mice (Figure S5). This contrasts with adult zebrafish *spock1^hm41/hm41^* mutants, which have a defective BBB (Figure 1), indicating species-specific differences in the adult BBB.

**Figure 4.**
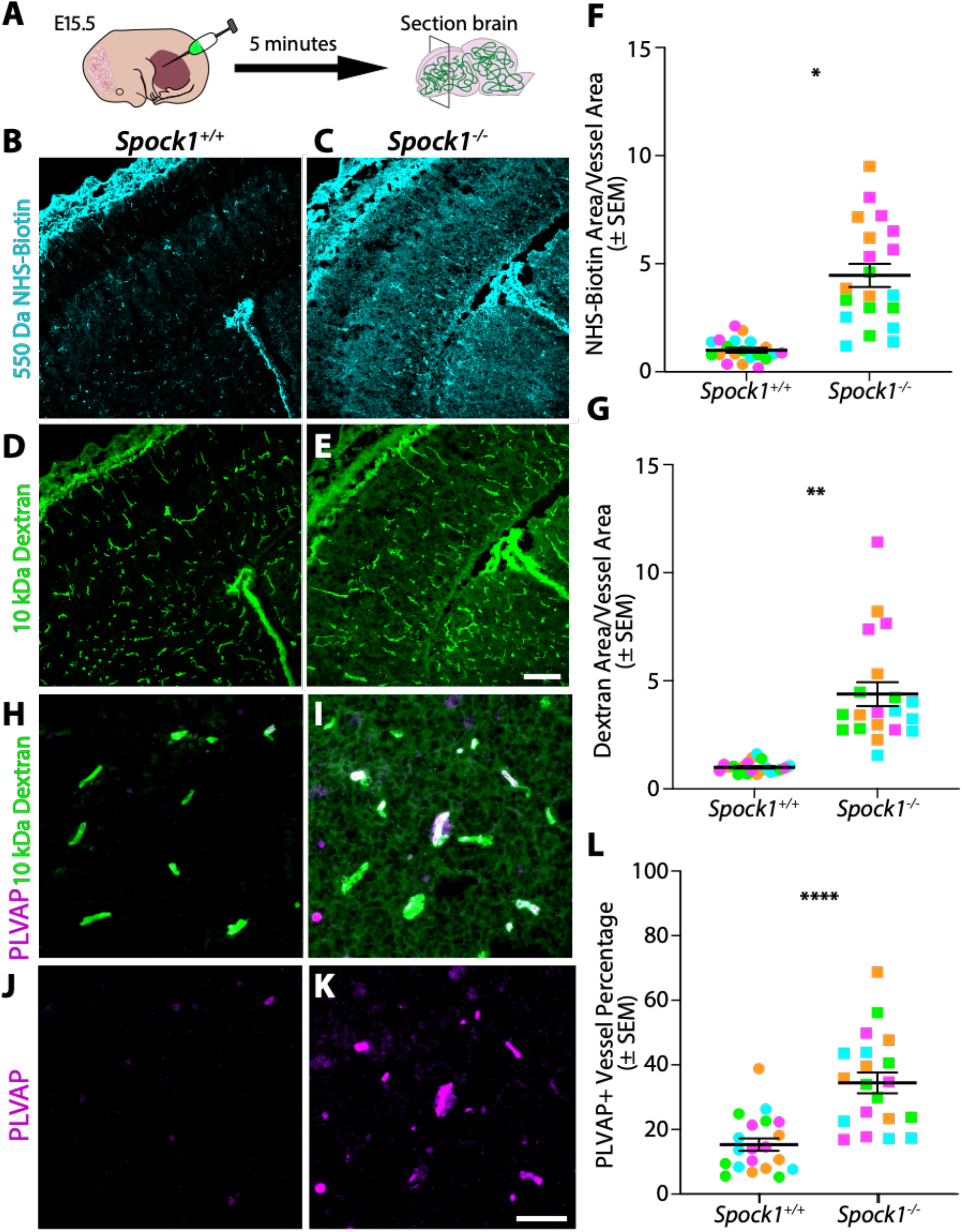
Spock1 plays a conserved role in inducing barrier function during embryonic development in mice. (A) Schematic of functional tracer leakage assays in embryonic day 15.5 (E15.5) mice. (**B-E**) Wild type mice confine both injected 550 Da NHS-Biotin (**B**) and 10 kDa Dextran (**D**) within the vasculature, as previously reported. However, *Spock1^-/-^* mice leak both NHS-Biotin (**C**) and Dextran (**E**) into the brain parenchyma. (**F-G**) Quantification of the total area of NHS-Biotin leakage (F) and Dextran leakage (G) normalized to vessel area where a ratio of 1 indicates no leakage reveals a significant increase in extravasation of both tracers in the *Spock1^- /-^* embryos (p=0.0213 (F) and p=0.0048 (G) by nested t test). (**H-K**) Zoomed in view of 10 kDa Dextran (green) confined within the vasculature of wild type (H) vessels and leaked out into the cortex of *Spock1^-/-^* knockouts (I). This increased BBB permeability in *Spock1^-/-^* mice is accompanied by an increase in PLVAP (magenta) expression in the vasculature (K). (**L**) Quantification of PLVAP expression within the vasculature reveals a significant increase in PLVAP expression in the *Spock1^-/-^* embryos (p<0.0001 by nested t test). N=4 embryos for each genotype, marked by unique colors, with 5 sections analyzed per embryo. Scale bars represent 100 µm (E) and 50 µm (I).

### Vascular cells are the only altered cell types in *spock1* mutant brains

To determine how Spock1, a neuronally produced and secreted proteoglycan, signals to and regulates brain endothelial cell BBB properties, we turned to scRNA-seq of dissected 5 dpf *spock1^hm41/hm41^* mutant and wild type brains, allowing us to globally assess cell type specific molecular changes occurring in the mutant brains. With Leiden clustering of the scRNA-seq data we defined cell clusters containing neurons, glial, and vascular cell types (Figure S6). We performed differential gene expression (DGE) analysis for each cluster and did not observe any changes in the neuronal or glial populations, but did observe significant changes in the vascular cluster (Table S2). When we subclustered the vascular population, we were able to resolve endothelial cells, pericytes and vascular smooth muscle cells (vSMCs; Figure 5A). Due to the low vascular cell numbers present in the scRNA-seq data, we prioritized candidate genes for subsequent validation by their fold-change in gene expression rather than statistical significance. These analyses suggested molecular changes in the mutant endothelial cells indicative of a leaky phenotype, with increased expression of *plvapb* and decreased expression of the tight junction protein *cldn5b* (Figure 5B, Table S3). Interestingly, *spock1* mutants had decreased expression of the melanoma cell adhesion molecule *mcamb* (also known as CD146) in both endothelial cells and pericytes (Figure 5B and 5C). This decreased expression in the vasculature was validated by HCR FISH, with minimal expression in mutant midbrain pericytes but normal levels in hindbrain pericytes, which maintain BBB function in *spock1^hm41/hm41^* mutants (Figure 5D-G). CD146 has previously been shown to be required for BBB integrity, as *CD146^-/-^* mice exhibit BBB breakdown due to decreased pericyte coverage and downstream loss of endothelial expression of Cldn5 (Chen et al., 2017).

**Figure 5.**
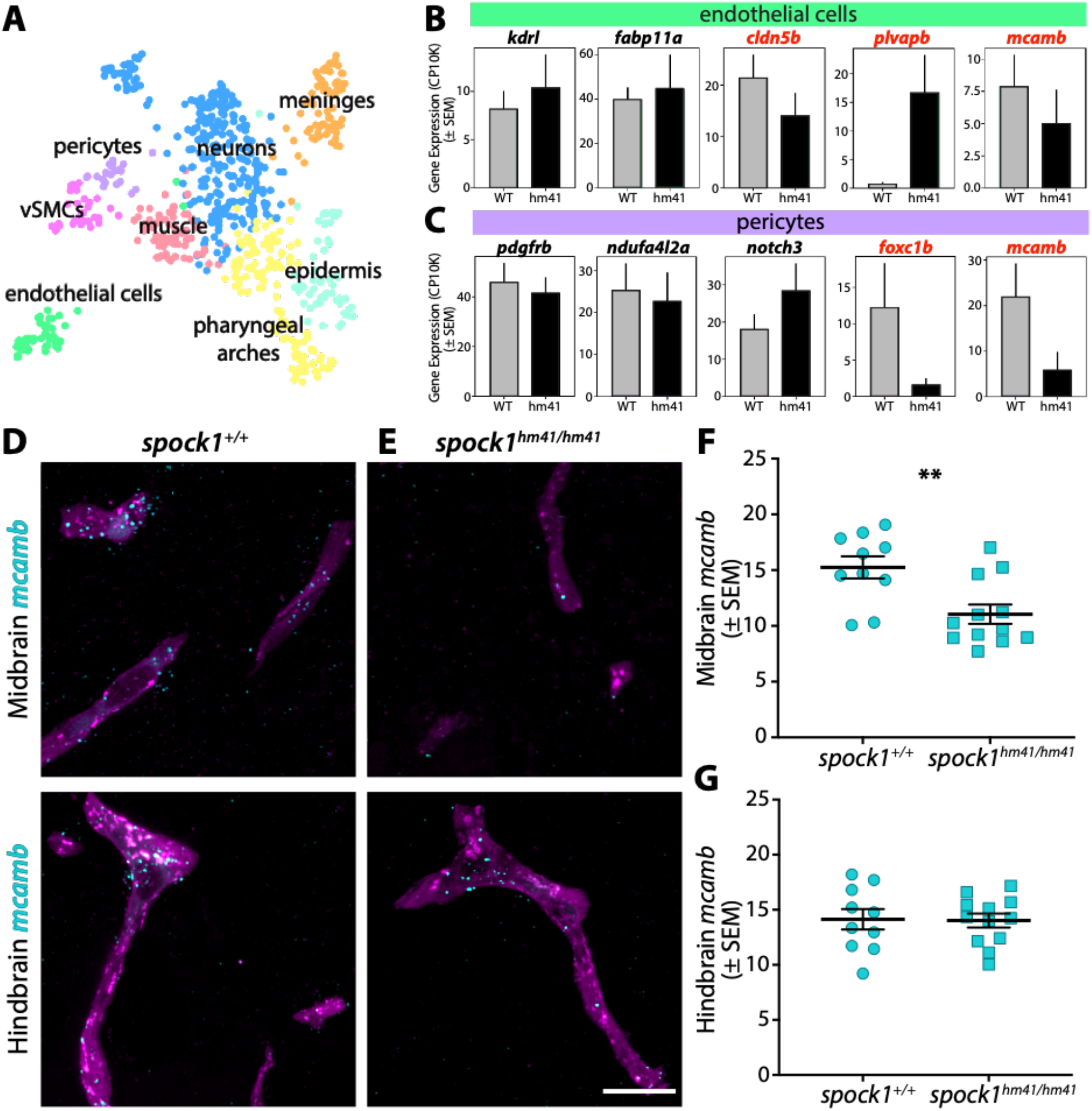
*Spock1^hm41/hm41^*mutant vascular cells have reduced expression of BBB regulators. (**A**) UMAP of the subclustered vascular cells separated by cell type, with pericytes (purple) separating from vascular smooth muscle cells (vSMCs, pink) and endothelial cells (green). (**B-C**) Mean gene expression in wild type (WT, grey bars) and *spock1* mutant (hm41, black bars) endothelial cells (B) and pericytes (C). Error bars represent SEM. Mutants appear to have lower levels of *mcamb* in both pericytes and endothelial cells. (**D-G**) HCR FISH reveals strong expression of *mcamb* (turquoise) in wild type vasculature (magenta) (D), both in the midbrain and hindbrain. *Spock1* mutants have significantly reduced expression of *mcamb* in the midbrain (E, F), but normal levels in the hindbrain (G), where no leakage is observed. Scale bar represents 10 µm. ** p=0.0044, **** p<0.0001 by unpaired t test.

### Exogenous SPOCK1 restores barrier function

To test the ability of Spock1 to rescue the mutant leakage phenotype, we injected recombinant human SPOCK1 (rSPOCK1) protein directly into the brain of *spock1^hm41/hm41^* mutant larvae at 5 dpf and assessed brain permeability at 6 dpf. While control animals injected with PBS maintained high levels of brain permeability (Figure 6A), larvae that received at least 2.3 ng of rSPOCK1 per mg fish body weight showed a 50% reduction in brain permeability following a single dose (Figure 6B and 6C). These results further confirm that SPOCK1 is able to act cell non-autonomously to rescue the mutant BBB leakage, as these injections were targeted broadly to the neural tissue rather than the endothelial cells, and demonstrate functional conservation of Spock1 from human to zebrafish.

**Figure 6.**
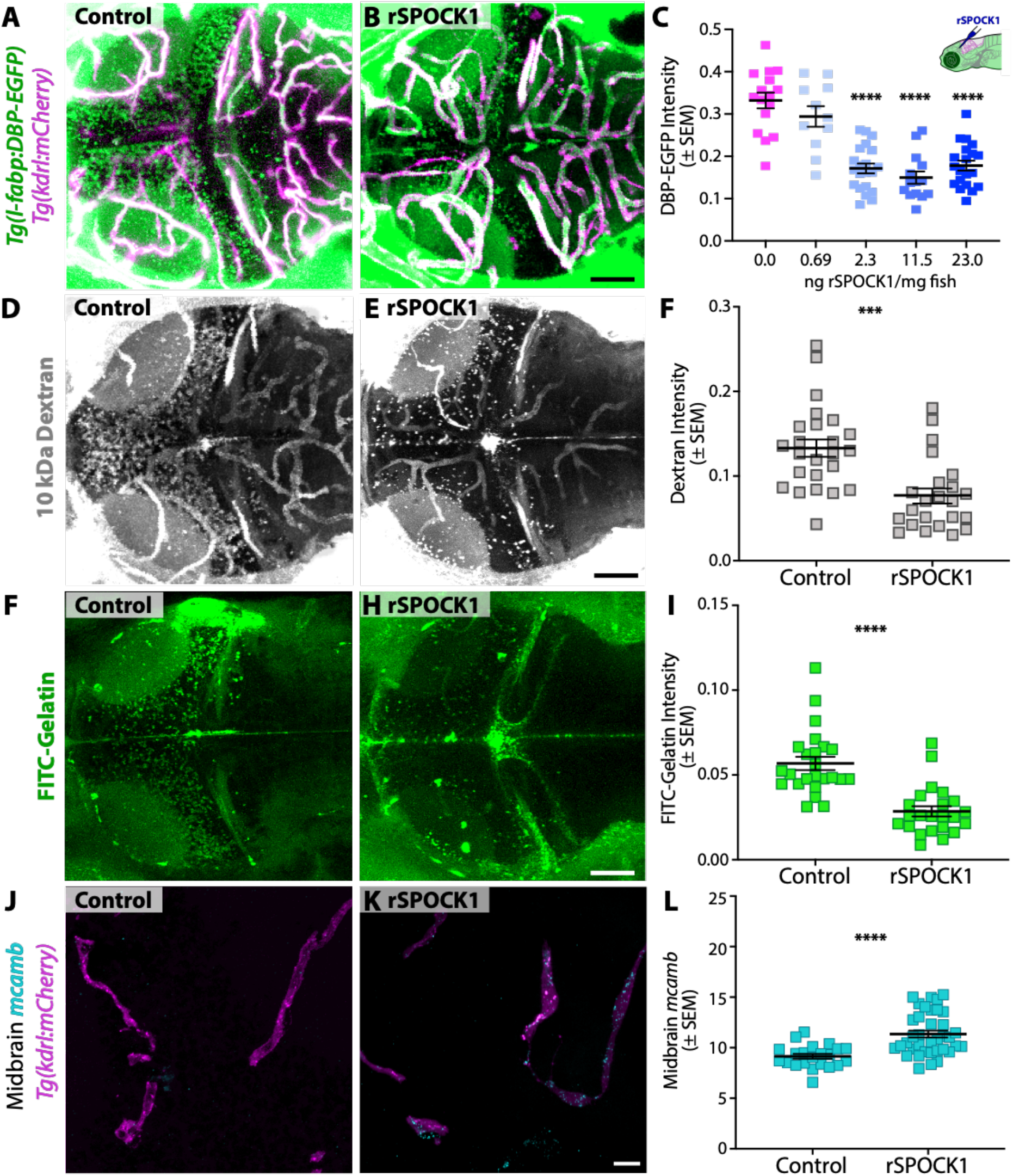
Spock1 induces BBB properties by modulating the brain microenvironment, altering vascular cell biology non-autonomously. (**A-C**) A single intracranial injection of human rSPOCK1 into the brain at 5 dpf reduces mutant leakage of DBP-EGFP at 6 dpf about 50% (B) compared to controls injected with PBS alone (A), quantified in C. (**D-F**) Similarly, injection of rSPOCK1 at 4 dpf reduces mutant leakage of 10 kDa Dextran at 5 dpf (E), quantified in F. (**F-I**) The addition of rSPOCK1 reduces gelatinase activity in the mutant brain (H) compared to control injected mutants (F), quantified in I. (**J-L**) Vascular expression of *mcamb* in the midbrain is partially restored following rSPOCK1 intracranial injections (K), quantified in L. Scale bars represent 50 µm (C, E, F) and 10 µm (K). **** p<0.0001 by 2way ANOVA compared to PBS injected control *spock1^hm41/hm41^* mutants in C and by t test in I and L, *** p<0.001 by t test in F.

To determine whether this restoration of barrier function corresponds to Spock1’s regulation of the MMP activity, we repeated the intracranial injections of 11.5 ng rSPOCK1/mg of fish in *spock1^hm41/hm41^*mutants at 4 dpf and assessed barrier function and gelatinase activity at 5 dpf. While 5 dpf control injected *spock1^hm41/hm41^* mutants continued to display high levels of BBB permeability and gelatinase activity, rSPOCK1 reduced mutant leakage of 10 kDa Dextran about 50% (Figure 6E and 6F), just as it did when we injected rSPOCK1 at 5 dpf and assessed BBB function at 6 dpf (Figure 6C). This restoration of BBB function was accompanied by a decrease in gelatinase activity (Figure 6H and 6I). Finally, to determine whether Spock1 lies upstream of vascular expression of the cell adhesion molecule *mcamb,* we assessed *mcamb* expression in the mutant midbrain vasculature at 5 dpf with or without the addition of rSPOCK1. While *spock1^hm41/hm41^* mutants injected with PBS displayed negligible levels of *mcamb* expression in the midbrain (Figure 6J), mutants injected with rSPOCK1 had a 50% increase in *mcamb* expression in the vasculature (Figure 6K and 6L), with a noticeable increase in *mcamb*+ pericytes. Interestingly this 50% increase in *mcamb* expression directly correlates with the 50% reduction in leakage and gelatinase activity following rSPOCK1 injections (Figure 6F and 6I).

## Discussion

Together these data show that Spock1 is a secreted, neuronally expressed extracellular signal that regulates BBB permeability in zebrafish and mouse without altering vascular patterning during development (Figures 1 and 4). Mechanistically, this work suggests a model whereby Spock1 regulates the brain extracellular environment via its inhibition of gelatinase activity, thereby providing the appropriate chemical or mechanical signals for vascular pericytes and endothelial cells to communicate properly with one another. This critical pericyte-endothelial interaction thereby prompts a BBB transcriptional program, including increased expression of tight junction (*cldn5b)* and cell adhesion (*mcamb*) molecules, decreased expression of leaky endothelial genes (*plvapb*), and reduced transcytosis. Thus, loss of Spock1 function contributes to increased BBB permeability by disruption of the complex extracellular scaffold that surrounds and supports the vasculature during development. Together this work reveals how a novel signal from the brain microenvironment can regulate the vasculature to give rise to the special properties of the BBB and provides new targets for its therapeutic modulation.

Despite the ubiquitous expression of *spock1* throughout the CNS, *spock1* loss of function zebrafish mutants display regional leakage in the forebrain and midbrain and a completely intact hindbrain barrier. Spock1 is not alone in having region-specific regulation of barrier properties despite pan-CNS expression. A recent study demonstrated that the pericyte-produced ECM molecule vitronectin is required for functional retinal and cerebellar barriers despite its expression by all CNS pericytes (Ayloo et al., 2022). Even mutants in canonical Wnt signaling have varying levels or regionalized angiogenic or barrier defects despite wide-spread sources of Wnt signals and receptors throughout the brain (Cho et al., 2017; Junge et al., 2009; Wang et al., 2012, 2018; Xu et al., 2004; Zhou and Nathans, 2014; Zhou et al., 2014). Interestingly, the regions with the highest susceptibility to single gene mutations in canonical Wnt signaling are the retina and the cerebellum, while the cortex can maintain a functional BBB even with mutations in several genes in the Wnt pathway (Wang et al., 2018). These data suggest that different regions of the brain are either molecularly or cellularly (or both) distinct, perhaps in nuanced ways that are often masked by current detection methods. For example, while the neurovascular unit of endothelial cells and pericytes in close contact with astrocytic endfeet appears uniform throughout the brain by immunostaining and EM, various scRNA-seq datasets have now shown that astrocytes are highly regionally distinct at a molecular level (Herrero-Navarro et al., 2021; Ohlig et al., 2021; Zhu et al., 2018) raising the possibility that they may respond to different signals. Pericytes in the brain, unlike in the rest of the body, arise from two discrete origins, mesodermal and neural crest (Ando et al., 2016; Trost et al., 2016). In zebrafish, the two origins have discrete domains, with the hindbrain being exclusively derived from mesodermal origins and the midbrain and forebrain being predominantly of neural crest origin (Ando et al., 2016) which interestingly matches the leakage boundaries in *spock1* mutants.

Here we demonstrate that Spock1 regulates brain endothelial cell identity to initiate the barrier program during development, both at molecular and subcellular resolution. However, this does not preclude the fact that other brain barriers may be similarly affected by loss of this signal. In addition to the BBB, the neural tissue is also protected by the blood-cerebrospinal fluid (CSF) barrier, which is created by the choroid plexus epithelial cells lining the brain ventricles (Wolburg and Paulus, 2010), and the arachnoid barrier, which is created by the epithelial cells beneath the dura that completely enwrap the brain (Abbott, 2004). These various brain barriers mature on different developmental timelines, with the blood-CSF barrier becoming impermeable by 4 dpf (Henson et al., 2014), then the BBB by 5 dpf (O’Brown et al., 2019), and the arachnoid barrier maturing after 9 dpf (Jeong et al., 2008) in zebrafish. While the focus of our study has been the BBB, it is possible that these additional brain barriers also play a role in the observed leakage of various tracers into the brain in *spock1* mutants.

The N-terminal region of Spock1 has previously been shown to regulate matrix metalloproteinase (MMP) activity, specifically gelatinases MMP-2 and 9 (Du et al., 2020). We show here that leaky *spock1^hm41/hm41^* mutants have increased gelatinase activity in the midbrain occurring in the same place as the observed decrease in the endothelial-pericyte basement membrane (Figure 3 and S1), suggesting that this excessive MMP activity may be responsible for the breakdown of the basement membrane. Although all the cells in the neurovascular unit secrete MMP-2 and 9, pericytes have recently been shown to produce MMP-9 in response to ischemia and TGF-β1 exposure, leading to increased barrier breakdown and neuronal damage (Takahashi et al., 2014; Underly et al., 2017). For this reason, many have proposed gelatinases as a therapeutic target for several neurological diseases that are accompanied with barrier breakdown including stroke, multiple sclerosis, Alzheimer’s diseases, and cerebral hemorrhage (Avolio et al., 2003; Kook et al., 2013; Rosell et al., 2008; Underly et al., 2017). Given the ability of a single dose of rSPOCK1 to mitigate much of the leakage in the mutants and restore proper vascular expression of BBB genes, we propose that Spock1 may serve as a new therapeutic target to reduce gelatinase activity in these neurological diseases and help restore barrier function.

## Acknowledgments

We thank members of the Megason laboratory for data discussion and comments on the manuscript; Dr. Zach O’Brown for discussions and comments on the manuscript; Ignas Mazelis for single cell demultiplexing software; the HMS Single Cell Core for single cell library preparation; Dr. Bela Anand-Apte (Cleveland Clinic) for providing the transgenic *l-fabp:DBP-EGFP* fish line (Xie et al., 2010) and Dr. Leonard Zon for providing the transgenic Tg(*kdrl:HRAS-mCherry*) line; and the HMS Electron Microscopy Core Facility, with special thanks to Louise Trakimas for all of her assistance in preparing the TEM samples. This work was supported by the Damon Runyon Cancer Foundation (N.M.O.), NIH R01HD096755 (S.G.M), an Allen Distinguished Investigator Award (C.G.), NIH R35NS116820 (C.G), and partially supported by Faculty Scholar grant from the Howard Hughes Medical Institute (C.G.). C.G. is an investigator of the Howard Hughes Medical Institute.

## Author Contributions

N.M.O., S.G.M. and C.G. conceived the project and designed experiments. N.M.O. performed all experiments and analyzed most data with the exception of the scRNA-seq which was performed in collaboration with and analyzed by N.B.P. and A.M.K. U.H. provided the *Spock1*^-/-^ mice. N.M.O., S.G.M. and C.G wrote the manuscript.

## Supplementary Materials

### Materials and Methods

#### Zebrafish Strains and Maintenance

Zebrafish were maintained at 28.5°C following standard protocols(Westerfield, 1993). All zebrafish work was approved by the Harvard Medical Area Standing Committee on Animals under protocol number IS00001263-3. Adult fish were maintained on a standard light-dark cycle from 8 am to 11 pm. Adult fish, age 3 months to 2 years, were crossed to produce embryos and larvae. For imaging live larvae, 0.003% phenylthiourea (PTU) was used beginning at 1 dpf to inhibit melanin production. These studies used the AB wild-type strains and the transgenic reporter strains Tg(*l-fabp:DBP-EGFP)*^Iri500^ (Xie et al., 2010), (Tg(*kdrl:HRAS-mCherry*)^s896^ (Chi et al., 2008), abbreviated as Tg(*kdrl:mCherry*) in the text, TgBAC(*pdgfrb:EGFP)*^ncv22Tg^ (Ando et al., 2016), abbreviated as Tg(*pdgfrb:EGFP*), and Tg(*plvapb:EGFP*)^sj3Tg^ (Umans et al., 2017), abbreviated as Tg(*plvapb:EGFP*) in the text.

#### *Spock1^hm41^* Mutants

*Spock1^hm41^* mutants were maintained in the double transgenic Tg(*l-fabp:DBP-EGFP; kdrl:mCherry)* background. Heterozygous fish were intercrossed for all leakage assays, with the exception of the time lapse microscopy experiments, where a heterozygous fish was crossed to a homozygous mutant, and the cell transplantation experiments, where homozygous mutants were in-crossed. All larvae were imaged prior to genotyping to identify wild type and mutant fish. The *spock1^hm41^* mutant line was genotyped using 5’-ACTGAGTGTTATTTTGTCATTGTGC-3’ and 5’-TGATGCTGATCTGAGAAGTTTAGCC-3’ primers followed by a HaeIII restriction digest, which does not digest the wild-type product (327 bp).

#### ***Spock1^hm43^*** Mutants

*Spock1^hm43^* mutants were generated and maintained in the double transgenic Tg(*plvap:EGFP; kdrl:mCherry)* background using CRISPR mutagenesis with two sgRNAs (5’- GTTCGTCACTTTAAAAGGAG-3’ and 5’-TCTCAGATGATGTTGAGTCT-3’) to target the *spock1* 5’ UTR and start codon. Heterozygous fish were intercrossed for all leakage assays. All larvae were imaged prior to genotyping to identify wild type and mutant fish. The *spock1^hm43^* mutant line was genotyped using 5’-TAAACTTCGATTGTGCTTTGGTTT G-3’ and 5’- AATGACTACAGACCTTTGAGGGAAC-3’ primers and KAPA HiFi Hotstart polymerase (Roche:KK2602) using a 58°C melting temperature. The mutant PCR product of 364 bp was easily distinguishable by gel electrophoresis from the wild type product at 935 bp.

#### Fluorescent Zebrafish Tracer Injections and Live Imaging

Larvae were immobilized with tricaine and placed in an agarose injection mold with their hearts facing upwards. 2.3 nl of Alexa Fluor 405 NHS Ester (Thermo Fisher: A30000) or Alexa Fluor 647 10 kDa Dextran (Thermo Fisher: D22914) fluorescently conjugated tracers (10 mg/ml) were injected into the cardiac sac using Nanoject II (Drummond Scientific, Broomall, PA). Larvae were then mounted with 1.5% low gelling agarose (Sigma: A9414) in embryo water on 0.17 mm coverslips and imaged live within 2 hours post injection on a Leica SP8 laser scanning confocal microscope using the same acquisition settings with 1 µm z-steps using a 25x water immersion objective. All quantification was performed on blinded image sets. For static images, parenchymal fluorescent tracer intensity was measured using ImageJ in the entire regional parenchyma outside of the vasculature in 60 µm thick maximum intensity projections of the larval brains. These projections began on average 15 µm below the mesencephalic vein to reduce the effects of potential leakage diffusion from the surface vessels and had the vasculature masked and removed for intensity quantification. These parenchymal tracer intensity values were then background subtracted and normalized to the tracer intensity within the vasculature to account for differential amounts of circulating tracer between fish. For time lapse imaging, Dextran intensity was measured in six parenchymal regions of average intensity projections of the time lapse videos and averaged as a single value per fish and similarly normalized to the average blood vessel luminal fluorescence intensity at each time point. Pericyte and vascular double transgenics were similarly imaged on the Leica SP8 for quantification of vascular defects. Additional images were taken on a Zeiss LSM 980 with Airyscan 2 with 0.21 µm z-steps using a 20x water immersion objective for increased cellular resolution, allowing us to detect gaps between the two cell types in vivo.

#### Linkage Mapping

Larvae from 2 separate crosses were screened for leakage of DBP-EGFP at 5 dpf and pooled into 3 groups of 5 to 6 leaky or wild type heads. RNA was extracted from the pools using RNeasy mini kit and ribo-depleted. RNA sequencing libraries were prepared using Wafergen Directional RNA-Seq kits and sequenced on NextSeq High-Output sequencers producing 75 bp paired end reads. Reads were mapped to the GRCz11 genome using tophat and bowtie2. Linkage mapping was performed on the mapped reads using RNAmapper(Miller et al., 2013) with the following specifications: zygosity=25, coverage=1, linkedRatio=0.96, neighbors=10. Differential gene expression analysis was performed on these libraries using rsem(Li and Dewey, 2011).

#### CRISPR Mutants

*Gstp2* crispant fish were generated by injection of Cas9 protein and 3 guide RNAs (5’- CAGCTGCCTAAATTTGAAGA-3’, 5’-GCGTTGGAAACTTACACATG-3’, and 5’-GTGAG AGTGTAGGGAGCCAC-3’) into 1-cell fertilized double transgenic Tg(*l-fabp:DBP-EGFP; kdrl:mCherry)* embryos. *Csf1ra* crispant fish were generated similarly with 4 guide RNAs (5’- CTGCTCACCAACAGCCGAG-3’, 5’-GTGTCTTCTGACCGACCCGG-3’, 5’-CTCGTC TTCATGCTTCACG-3’, and 5’-AGTGACACCTTCTCCATGG-3’), as were *spock1* crispants (5’- GTAGCCGACAGAAAGAGAGG-3’, 5’-GAGTCGCAGGAGTTGAACAG-3’, 5’- GACAGTGAACCTTCATGCAG-3’, and 5’-TGTCCGGGCAGGCAAGGGCA-3’). F0 crispants were analyzed for leakage of the DBP-EGFP tracer outside of the kdrl:mCherry labeled vasculature at 5 dpf.

#### HCR Fluorescent In Situ Hybridization (FISH)

*Slide*: HCR RNA in situ hybridization (Molecular Instruments) experiments on 14 µm cryosections of fixed 5 dpf larvae were performed as previously described (Choi et al., 2018; O’Brown et al., 2019). Briefly, sections were air dried and re-fixed in 4% paraformaldehyde (PFA) in PBS for 10 minutes at room temperature. Following fixation, slides were washed in PBS and then permeabilized using 1 µg/ml Proteinase K (ThermoFisher) for 5 minutes, followed by PBS washes and refixation. Tissues were further permeabilized by an ethanol dehydration series of 50%, 70% and two rounds of 100% ethanol and air dried for 5 minutes. Dried slides were put into probe hybridization solution for at least 10 minutes at 37°C. Subsequently, the probes for *spock1, kdrl, elavl3,* and *mcamb* were added to the slides at a final concentration of 4 nM to hybridize overnight at 37°C. The next day, slides were washed with a series of wash buffer to 5x SSCT (5x SSC with 0.1% Tween 20). Excess liquid was then removed and samples were immersed in amplification buffer at room temperature for 30 minutes prior to hairpin amplification, which occurred overnight at room temperature. Slides were then washed in 5x SSCT, washed in 5x SSC and mounted with Fluoromount-G (Electron Microscopy Sciences).

*Whole mount*: Larvae were fixed in 4% PFA overnight and then stored in methanol at −#x00B0;C until staining was performed. All staining was performed in PCR strip tubes. Larvae were rehydrated in a series of methanol/PBST (0.1% Tween) washes and then permeabilized with 30 µg/ml Proteinase K for 45 minutes at room temperature. Larvae were then refixed in 4% PFA for 20 minutes and treated with probe hybridization buffer at 37°C. Subsequently, the *spock1* probe was added at a final concentration of 4 nM to hybridize overnight rocking at 37°C. The next day, larvae were washed with a series of wash buffer to 5x SSCT (5x SSC with 0.1% Tween 20). Excess liquid was then removed and samples were immersed in amplification buffer at room temperature for 30 minutes prior to hairpin amplification, which occurred overnight rocking at room temperature. Larvae were then washed and imaged in 5x SSCT.

Images for quantification were collected on a Leica SP8 laser scanning confocal microscope using the same acquisition settings with 0.2 µm z-steps using a 25x water immersion objective or a 63x oil immersion lens. All quantification was performed on blinded image sets of sections from a single slide treated with all of the same reagents and imaged on the same day. To measure levels of *mcamb* expression, we used ImageJ to first subtract tissue autofluorescence (captured in the 405 channel) from the *mcamb* expression in maximum intensity projections and then the vasculature was manually traced and average intensity was measured. Additional images were taken on a Zeiss LSM 980 with Airyscan 2 with 0.21 µm z-steps using either a 20x or a 40x water immersion objective to further enhance the resolution of the *mcamb* signal.

#### Transplantation

Donor embryos were injected with 2.3 nl of 10 mg/ml Alexa Fluor 647 10 kDa Dextran (Thermo Fisher: D22914) at the 1-cell stage to distinguish them from host cells. Following injection, embryos were incubated at 28.5°C until transplantation. Host and donor embryos were dechorinated with 1 mg/ml Pronase (Roche:11459643001) at oblong stage and transferred to transplantation agarose dishes in 1/3 Ringer’s buffer. Unfertilized or injured embryos were discarded. To generate clonal sources secreting wild type or mutant Spock1, approximately 40– 80 cells were transplanted from sphere stage dextran labeled donor embryos into sphere stage wild type and mutant hosts (similar to previous studies(Pomreinke et al., 2017)). Embryos recovered overnight in 1/3 Ringer’s buffer and were subsequently transferred to Danieau Buffer with PTU. Tracer injections were performed at 5 dpf, as described above, with 1 kDa Alexa Flour 405 NHS Ester. Blinded image sets were analyzed using ImageJ. Individual larval brains were segmented into 10 µm thick maximum intensity projections spanning the entire brain. Average NHS tracer intensity was measured in 10 µm wide swaths connecting donor cells to the nearest blood vessel and normalized to average blood vessel tracer intensity. The median tracer intensity for a given distance (D) to the nearest donor cell was calculated for each individual fish.

#### Transmission Electron Microscopy (TEM)

Larvae (7 dpf) were anesthetized with tricaine and injected with 2.3 nl of 5 nm NHS-activated gold nanoparticles (Cytodiagnostics: CGN5K-5-1, ∼1.1^14^ particles/ml in PBS) just as for the fluorescent tracer injections. After 5 minutes of circulation, the larvae were initially fixed by immersion in 4% paraformaldehyde (VWR:15713-S) /0.1M sodium-cacodylate (VWR:11653). Following this initial fixation, larvae were further fixed for 7 days in 2% glutaraldehyde (Electron Microscopy Sciences: 16320)/ 4% paraformaldehyde/ 0.1M sodium-cacodylate at room temperature. Following fixation, larvae were washed overnight in 0.1M sodium-cacodylate. Entire larval heads were post-fixed in 1% osmium tetroxide and 1.5% potassium ferrocyanide, dehydrated, and embedded in epoxy resin. Ultrathin sections of 80 nm were then cut from the block surface and collected on copper grids. Grids were imaged using a 1200EX electron microscope (JEOL) equipped with a 2k CCD digital camera (AMT) and quantified using ImageJ (NIH). Vesicular density values were calculated from the number of non-clathrin coated small vesicles less than 100 nm in diameter or large vesicles greater than 200 nm in diameter per µm^2^ of endothelial area for each image collected. Average pericyte basement membrane (BM) thickness was quantified by measuring the total BM area divided by the length of the pericyte-endothelial contact. All images for analysis were collected at 12000x magnification. 10-15 vessels were quantified for each fish, with each color representing a different fish.

#### Intracranial rSPOCK1 Injections

Larvae were anesthetized with tricaine and placed in an agarose injection mold with their heads facing upwards. 2.3 nl of recombinant human SPOCK1 (rSPOCK1; R&D Systems: 2327-PI-050) at a range of 0.001-10 mg/ml was injected directly into the midbrain using Nanoject II (Drummond Scientific, Broomall, PA). Following injections, larvae recovered in normal embryo water for 1 day. The effects of rSPOCK1 were either assess in live fish using functional tracer leakage assays and in vivo gelatin zymography or in fixed samples followed by slide HCR in situ hybridization, as described above.

#### In vivo Gelatinase Zymography

Larvae were anesthetized and intracranially injected as for rSPOCK1 with 2.3 nl of FITC-Gelatin (Biovision: M1303) at 5 mg/ml. After one hour, larvae were also injected with 10 kDa Dextran in the cardiac sac to assess BBB function and then mounted with 1.5% low gelling agarose (Sigma: A9414) in embryo water on 0.17 mm coverslips and imaged live within 2 hours post injection on a Leica SP8 laser scanning confocal microscope using the same acquisition settings with 1 µm z-steps using a 25x water immersion objective. All quantification was performed on blinded image sets. Parenchymal FITC-Gelatin probe intensity was measured using ImageJ in the entire midbrain and normalized to the average intensity in the ventricles, which have high gelatinase activity, to account for fish to fish injection variation. Regions of obvious damage at the injection site were not quantified.

#### Mouse Maintenance

*Spock1*^-/-^ knockout mice (Röll et al., 2006) were obtained from Ursula Hartmann’s lab and were backcrossed to and maintained on a C57Bl/6 background. Mice were genotyped with primers 5’- GCCACTGGTCATTGTCTAGG-3’, 5’- TGTGCCCAGTCATAGCCGAATAGC CTCTCC-3’, and 5’- GCTTGAGGTAGCCCTGTTGTCACC-3’ using KAPA HiFi Hotstart polymerase (Roche:KK2602). This PCR reaction produced a 185 bp wild type band and a 750 bp knockout band in heterozygous animals. All animals were treated according to institutional and US National Institutes of Health (NIH) guidelines approved by the Institutional Animal Care and Use Committee (IACUC) at Harvard Medical School under protocol IS00000045-6.

#### Mouse Tracer Injections

Heterozygous *Spock1*^+/-^ mice were intercrossed and used for tracer leakage assays performed at embryonic day 15.5 (E15.5) as previously described(Ben-Zvi et al., 2014). In brief, 5 µl of tracer cocktail (10 mg/ml EZ-Link NHS-Biotin (Thermo Fisher: 20217) and 5 mg/ml 10 kDa Dextran Alexa Fluor 488 (Thermo Fisher: D22910)) was injected into the liver of each embryo and allowed to circulate for 5 minutes. Embryonic heads were fixed by immersion in 4% paraformaldehyde overnight at 4°C and then frozen in TissueTek OCT (Sakura). 20 µm thick sections were then collected and immunostained with Streptavidin Alexa Fluor 405 (1:200; Thermo Fisher: S32351) and rat anti-PLVAP (1:200; BD Biosciences:553849). All embryos were injected and fixed for processing blind before genotyping. Adult mice were retroorbitally injected with 0.5 mg of EZ-Link NHS-Biotin per g of mouse body weight. The NHS-Biotin circulated for 30 minutes prior to brain dissections and fixation with 4% PFA. 30 µm thick cryosections were then collected and immunostained with Streptavidin Alexa Fluor 568 (1:200; Thermo Fisher: S32351) and goat anti-CD31 (1:100; R&D Systems: AF3628).

#### Single-cell RNA-sequencing

Larval zebrafish brains were dissected and split along the midbrain-hindbrain boundary in DMEM. Brains were dissociated using a modified protocol(Bresciani et al., 2018). Briefly, chemical dissociations were performed at 30.5°C using a mixture of 0.25% Trypsin-EDTA, Collagenase/Dispase (8 mg/mL) and DNaseI (20 µg/mL) for 15-20 minutes with gentle pipetting every few minutes and quenched with 10% fetal bovine serum in DMEM. Samples were hashed using Multi-seq as previously described with slight modifications(McGinnis et al., 2019). For each sample, 80 pmoles of lipid modified oligos (LMOs) were used to hash every 500k cells. The hashing reaction was quenched using 1% BSA in PBS and barcoded samples were subsequently pooled and washed with 1% BSA. The pooled cell mixture was resuspended in PBS + 0.1%BSA + 18% Optiprep at a final concentration of ∼300k cells/mL prior to single-cell capture with inDrops. The Single-cell Core (SCC) at Harvard Medical School captured single-cell transcriptomes and prepared NGS libraries as previously described(Klein et al., 2015) with a target capture of 45k cells per experiment. A summary of dissected tissues and corresponding sequencing information are described in Table S4.

Gene expression and hashtag libraries were mixed (9:1 ratio) and sequenced on an Illumina Nova-seq 6000 with the NovaSeq S2 kit. Reads were mapped onto the Zebrafish GRCz11 Release 101 genome assembly using previously described methods(Wagner et al., 2018). Hashtags were identified using custom code available on: https://github.com/AllonKleinLab/paper-data/tree/master/OBrown2021_ZebrafishBBB

Transcriptomes with greater than 350 UMIs were further filtered for viability by removing cells with >20% mitochondrial reads. Cell demultiplexing was performed manually by applying thresholds to delineate single cells from background and multiple populations. The resulting counts matrix was normalized to the mean UMIs per cell in the dataset.

To visualize the data, we first mean-normalized the data and identified highly variable genes from the wild type AB and RNF datasets (minimum of 3 transcripts per cell, minimum of 3 cells expressing gene, minimum V-score percentile of 85%). We then Z-scored counts for each gene and performed principal component analysis (PCA) with 50 components. Cells from the spock1 mutant libraries were projected into the same principal component subspace. A k-nearest neighbor graph (k=10) was generated based on the Euclidean distance in gene expression between cells within this subspace. The Leiden algorithm was used to cluster cells into subgroups(Traag et al., 2019). The neighborhood graph was embedded and visualized using UMAP and the data was explored interactively with SPRING(Weinreb et al., 2017) to aid in cell cluster annotation.

Cells belonging to the liver, pharyngeal arches, skin, and muscle were removed prior to differential gene expression (DGE) analysis. The filtered dataset was reanalyzed using the methods described above. The Wilcoxon rank-sum test was used to generate a list of potential genes that are differentially regulated (fold change > 2) between genotypes across each Leiden cluster (Table S2). Genes were only considered for DGE analysis if they were expressed in at least 5% of cells in a given cluster and had a minimum mean of 10 transcripts per cell. Cells belonging to the AB and RNF background were treated as a single wild type genotype for the analysis. The vascular cluster (Leiden 13) was subclustered and similarly analyzed for DGE (Table S3).

#### Data availability

The raw reads from the counts matrix and associated metadata from the single-cell experiments have been deposited to GEO.

All code for the single-cell analysis is available as interactive Jupyter notebooks here: https://github.com/AllonKleinLab/paper-data/tree/master/OBrown2021_ZebrafishBBB

Interactive exploration of the full single-cell data can be found here: https://kleintools.hms.harvard.edu/tools/springViewer_1_6_dev.html?client_datasets/2021_OBrown/2021_OBrown

Exploration of the vascular subcluster cells can be found here: https://kleintools.hms.harvard.edu/tools/springViewer_1_6_dev.html?client_datasets/2021_OBrown_Vasculature/2021_OBrown_Vasculature

## Supplemental Figures

**Figure S1.**
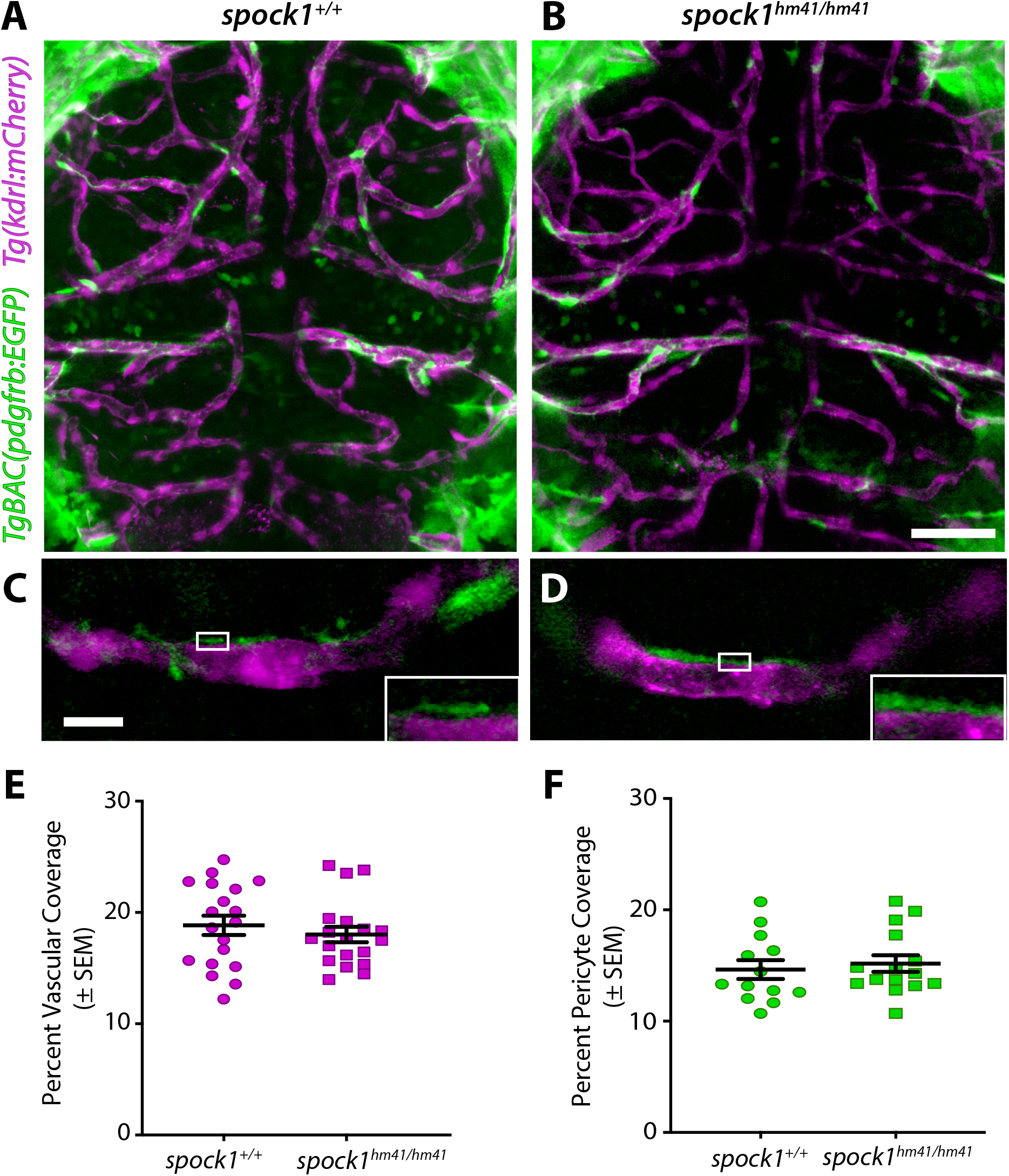
*Spock1^hm41/hm41^* mutants have normal vascular patterning. (**A-B**) Dorsal maximum intensity projection of 5 dpf wild type (**A**) and mutant (**B**) brains with endothelial cells marked by the kdrl:mCherry transgene (magenta) and pericytes marked by the pdgfrb:EGFP transgene (green). (**C-D**) Zoomed in view of vessels reveals diminished basement membrane between pericytes (green) and vessels (magenta) in mutants (**D**) compared to wild type siblings (**C**). This is further highlighted in the higher magnification insets, with a visible gap present in the wild type vessel (**C**) but not in the mutants (**D**). (**E-F**) Quantification of total vascular coverage of the brain (kdrl:mCherry+ area/total brain area, **E**) and total pericyte coverage of the vasculature (pdgfrb:EGFP+ area/kdrl:mCherry+ area, **F**), with each individual fish marked by a single point. Scale bars represent 50 µm (**B**) and 10 µm (**C**). T-test comparison reveals no significant difference between wild type and mutant fish for either vascular or pericyte coverage.

**Figure S2.**
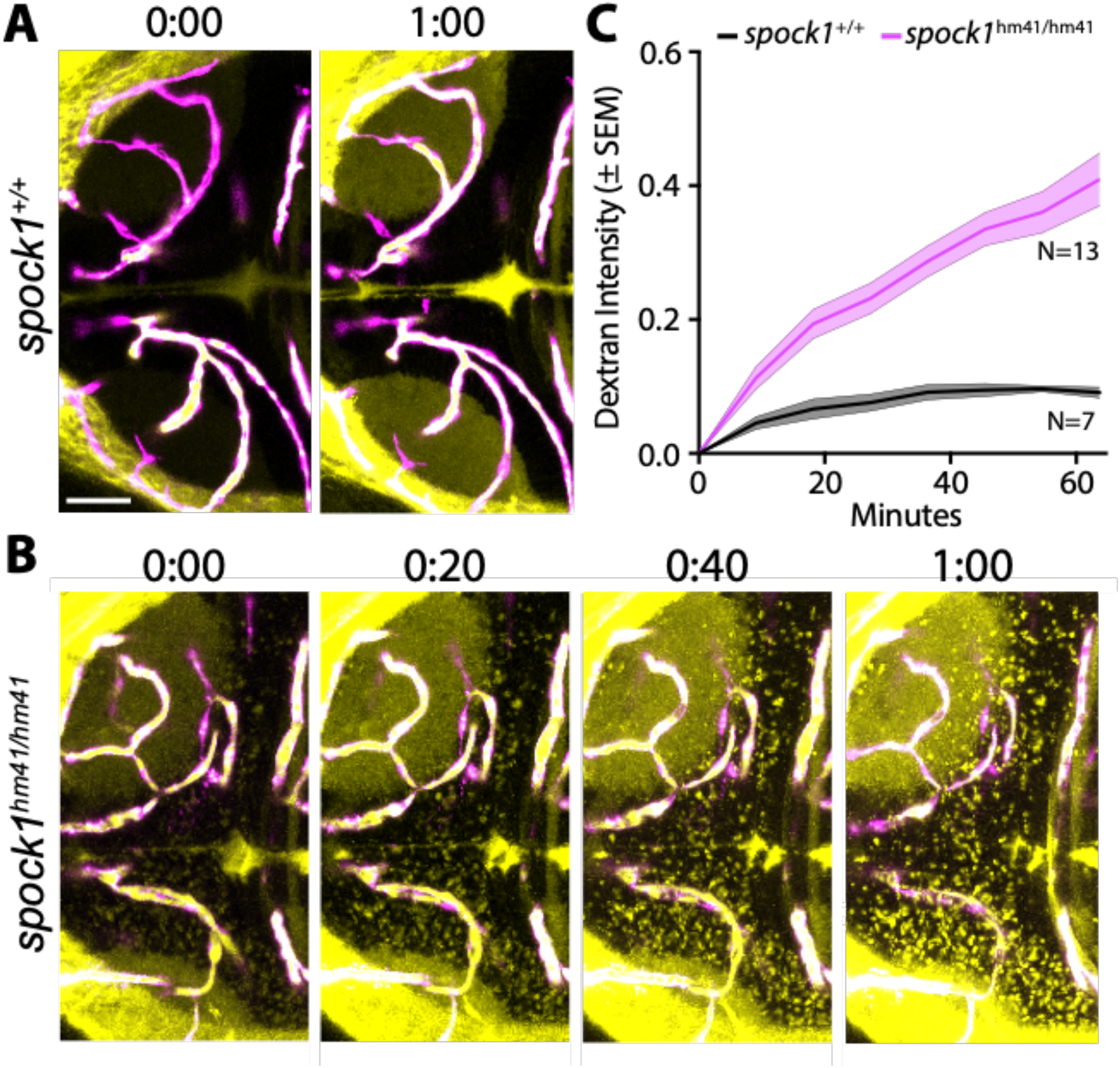
Time lapse imaging reveals leakage dynamics in *spock1^hm41/hm41^* mutants. (**A-B**) Dorsal maximum intensity projection of a wild type (A) and *spock1^hm41/hm41^* mutant (B) midbrain reveals steady accumulation of 10 kDa Dextran (yellow) outside of the vasculature (magenta) over the course of one hour in *spock1* mutants. (**C**) Quantification of Dextran accumulation in the midbrain parenchyma outside of the vasculature over time in wild type fish (black line) and *spock1^hm41/hm41^* mutants (magenta line). Scale bar represents 50 µm.

**Figure S3.**
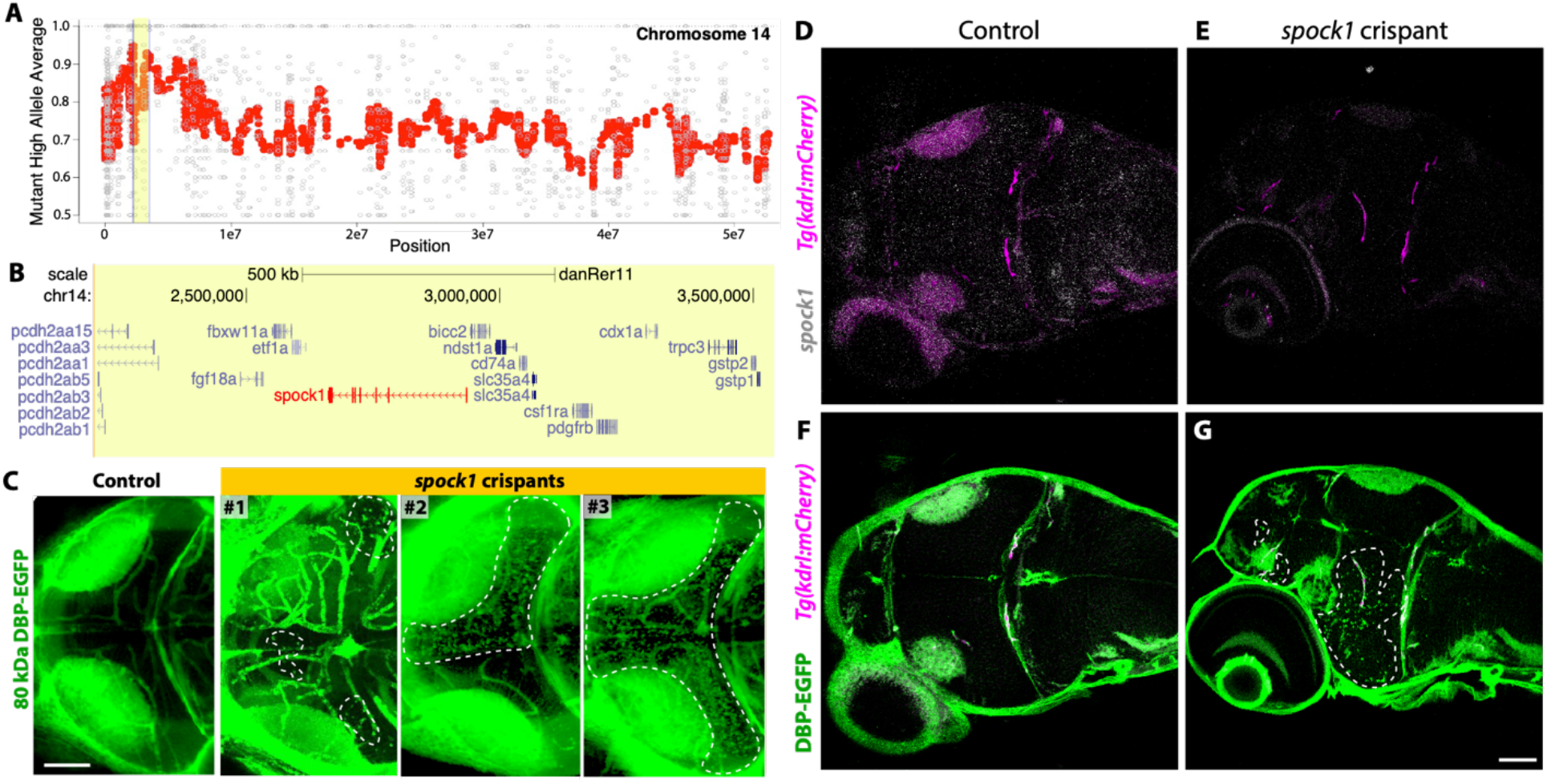
Leaky phenotype maps to *spock1* gene on chr14:2205271-3513919. (**A**) Manhattan plot of linkage on chromosome 14 with the average mutant alleles per 20 neighbors plotted in red. The region of highest linkage is highlighted in yellow. (**B**) Genome browser view of the highest linked region to the leaky phenotype. Eight of the genes within this region were expressed in the 5 dpf bulk RNAseq data: *fgf18a, spock1, bicc2, csf1ra, ndst1a, slc35a4, gstp1* and *gstp2*. Two of these genes were differentially expressed in leaky mutants compared to wild type (*csf1ra* and *gstp2*) and *spock1* (marked in red) had several SNPs that completely segregated in the leaky fish. (**C**) Mosaic *spock1* crispants display increased leakage of the transgenic DBP-EGFP tracer outside of the vasculature (outlined by white dashed line). (**D-E**) Dorsal view of whole mount HCR in situ for *spock1* (white) reveals expression throughout the brain of uninjected controls (D) but not in *spock1* crispants (E). (**F-G**) While controls confine the DBP-EGFP tracer (green) within the blood vessels (magenta) in the brain (F), *spock1* crispants display increased leakage into the brain parenchyma (G, outlined by white dashed line), corresponding with the loss of *spock1* expression (E). Scale bars represent 50 µm.

**Figure S4.**
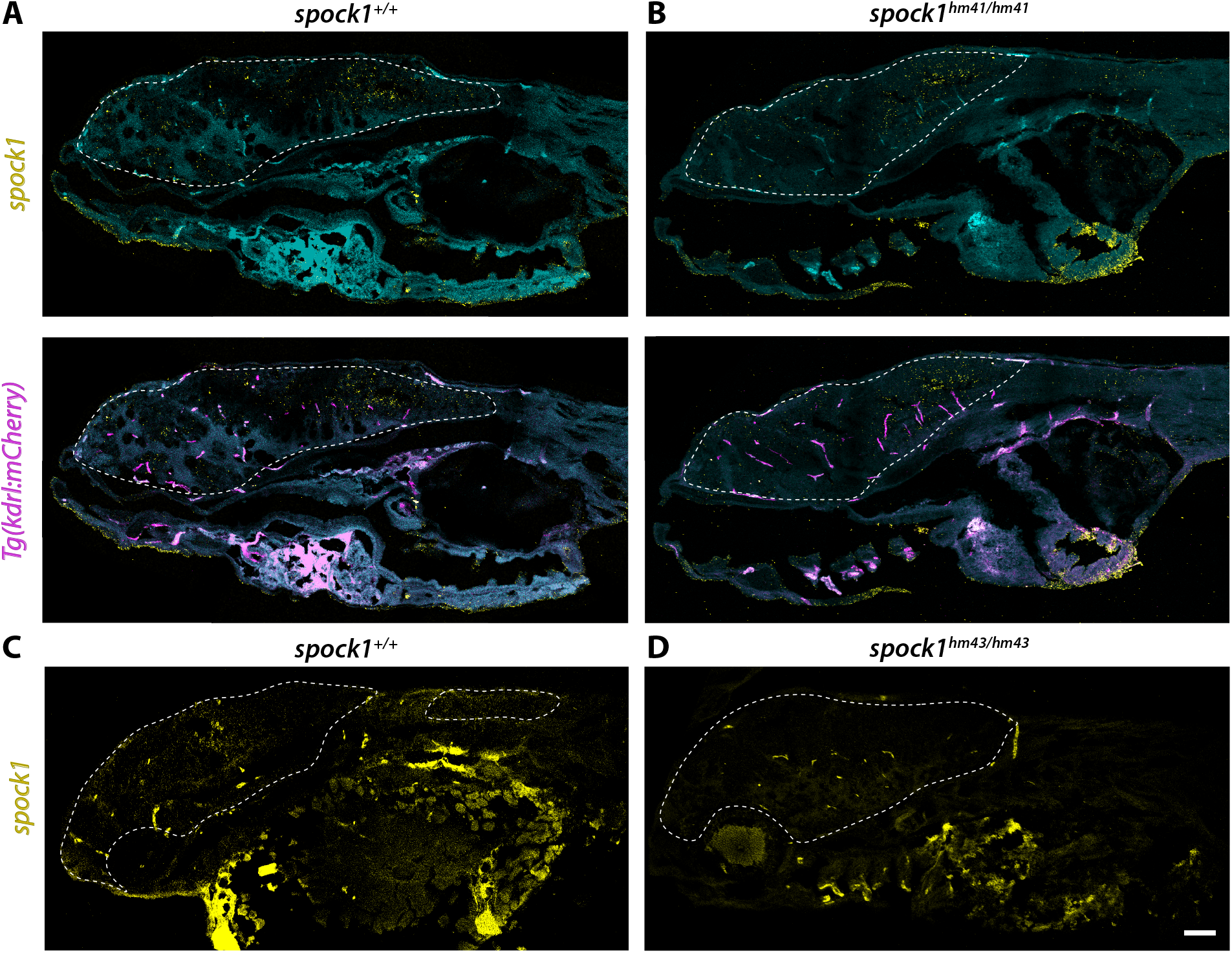
Expression of *spock1* is unaltered in *spock1^hm41/hm41^* mutants and absent in *spock1^hm43/hm43^*mutants. (**A-B**) *Spock1* expression (yellow) is found throughout the brain and spinal cord (outlined by a white dashed line) in wild type (A) and *spock1^hm41/hm41^* mutant fish (B) at 5 dpf. *Spock1* expression never colocalizes with vascular *Tg(kdrl:mCherry)* expression (magenta). Tissue autofluorescence is depicted in turquoise. (**C-D**) Unlike *spock1^hm41/hm41^* fish and wild type fish (C), *spock1^hm43/hm43^* mutants, which completely lack the 5’ UTR and start codon of *spock1*, do not have any detectable *spock1* expression in the brain (D). The remaining signal in the yolk and blood vessels is due to tissue autofluorescence. Scale bar represents 50 µm.

**Figure S5.**
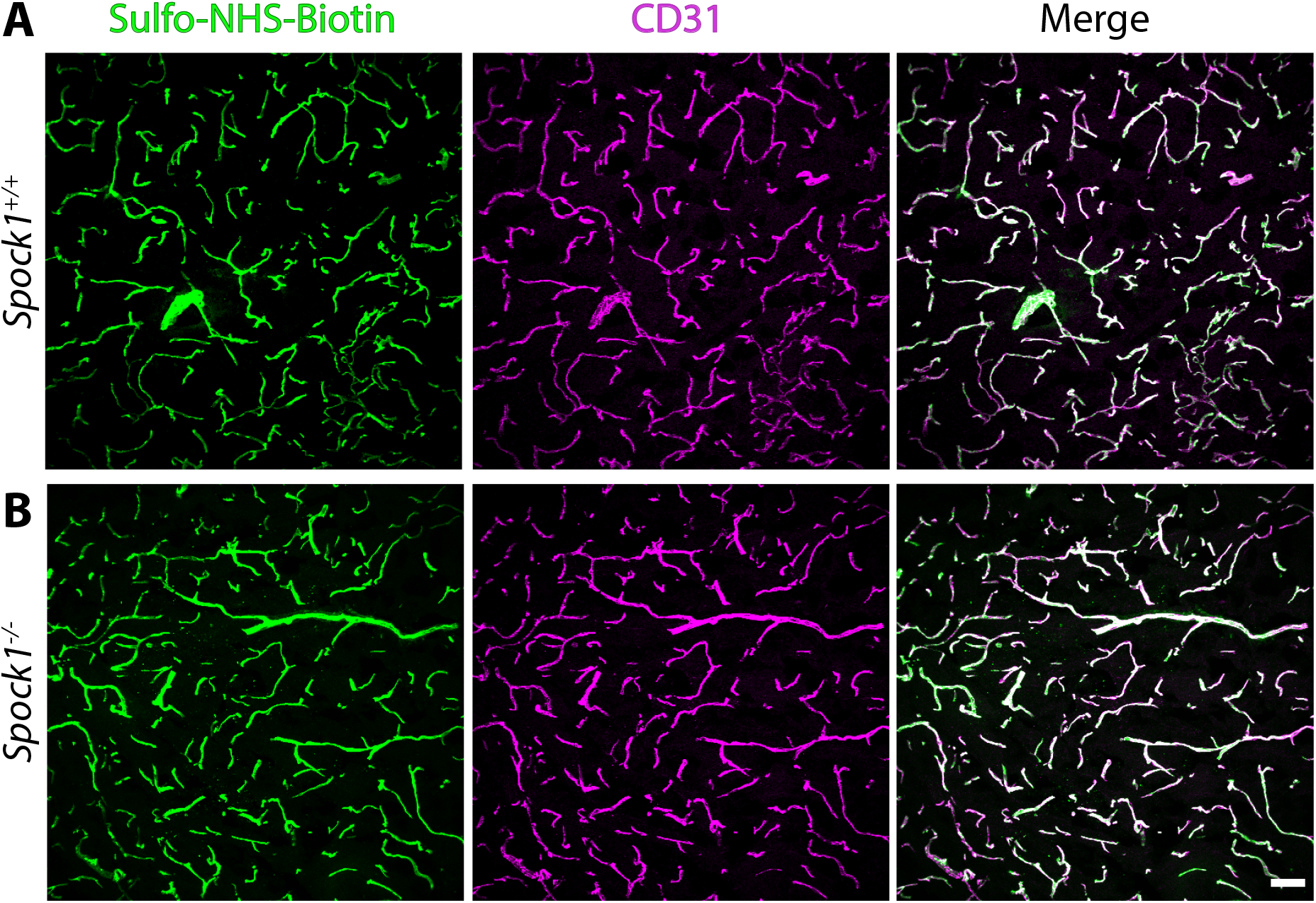
Adult *Spock1^-/-^*mice recover BBB function. (**A-B**) Tracer leakage assays in adult wild type (A) and *Spock1^-/-^* mice (B) reveals that both genotypes confine the injected Sulfo-NHS-Biotin tracer (green) within the CD31+ vasculature (magenta), indicating a functional BBB. Scale bar represents 50 µm.

**Figure S6.**
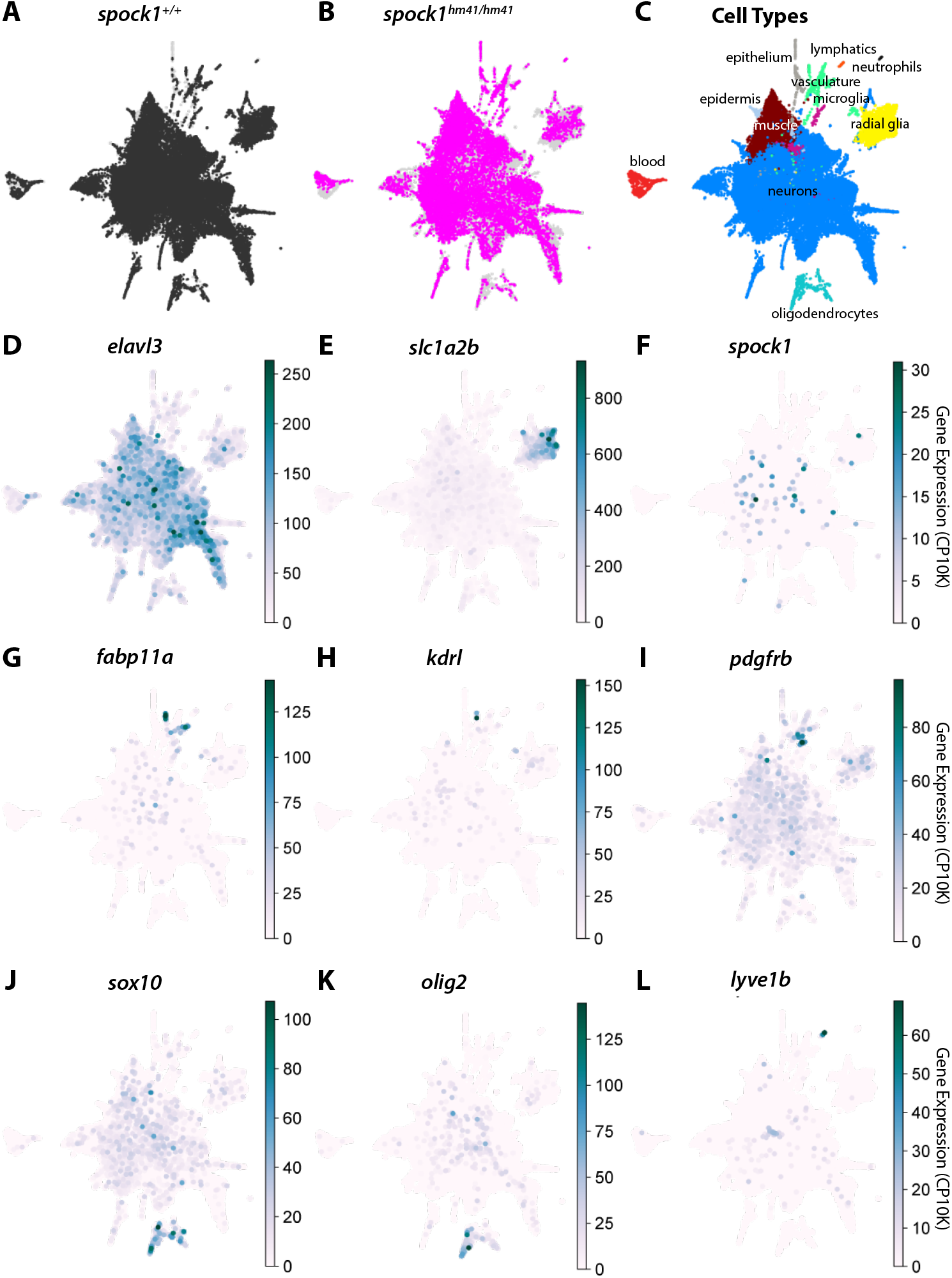
scRNA-seq captures all neurovascular cells in both mutant and wild type larvae. (**A-B**) UMAP of the whole brain dataset separated shows overlap between wild type (A) and *spock1^hm41/hm41^* mutant (B) cell type coverage, indicating that no cell type is absent in the mutant background. (**C**) UMAP of the total data set separated by annotated cell type, with the vast majority of sequenced cells being neuronal (blue). Our data does capture blood cells (red), microglia (magenta), oligodendrocytes (aqua), radial glia (yellow), and vascular cells (green). (**D-L**) Gene expression plots for *elavl3* (D) for neurons, *slc1a2b* (E) for radial glia, *fabp11a* (G) and *kdrl* (H) for endothelial cells, *pdgfrb* (I) for pericytes and neurons, *sox10* (J) and *olig2* (K) for oligodendrocytes, and *lyve1b* (L) for lymphatic endothelial cells reveals that *spock1* (F) is primarily expressed by neurons and absent from the vascular cells.

